# Organoid Pharmacotyping of Pancreatic Cancer Enables Functional Precision Oncology and Drug Repurposing

**DOI:** 10.64898/2025.12.01.691211

**Authors:** He Dong, Frederick S. Vizeacoumar, Yue Zhang, Nicholas Jette, Jared D.W. Price, Vincent Maranda, Lihui Gong, Tanya Freywald, Jeff Patrick Vizeacoumar, Mary Lazell-Wright, Saruul Uuganbayar, Jaylynn Quail, Alain Morejon Morales, Ayaan Rizvi, Rani Kanthan, Yuliang Wu, Anand Krishnan, Kathleen Felton, Bilal Marwa, Laura Hopkins, Gary Groot, John Shaw, Gavin Beck, Yigang Luo, Maurice Ogaick, Mike Moser, Andrew Freywald, Shahid Ahmed, Adnan Zaidi, Franco J. Vizeacoumar

**Affiliations:** Department of Oncology, College of Medicine, University of Saskatchewan, Saskatoon, S7N 5E5 Canada; Department of Pathology, College of Medicine, University of Saskatchewan, Saskatoon, S7N 5E5 Canada; Department of Surgery, University of Saskatchewan, Saskatoon, Canada; Department of Biochemistry, Microbiology and Immunology, University of Saskatchewan, Saskatoon, SK S7N 5E5, Canada; Department of Anatomy, Physiology, and Pharmacology, University of Saskatchewan, and Cameco MS Neuroscience Research Centre, 701 Queen St., Saskatoon, S7K 0M7, SK, Canada; Department of Pediatrics, Division of Pediatric Hematology/Oncology, Jim Pattison Children’s Hospital, Saskatoon, SK, Canada; Cancer Research Department, Saskatchewan Cancer Agency, 107 Wiggins Road, Saskatoon, S7N 5E5, Canada

## Abstract

**Purpose:** Pancreatic ductal adenocarcinoma (PDAC) remains one of the deadliest malignancies, with limited benefit from current cytotoxic regimens and poor predictive value of genomics alone. Patient-derived organoids (PDOs) represent a promising platform for functional precision oncology, yet systematic pharmacotyping of genomically annotated PDAC PDOs remains sparse.

**Experimental Design:** We established a clinically annotated panel of ten treatment-naïve PDAC PDOs spanning well-, moderately-, and poorly differentiated tumors. PDOs were evaluated for morphologic and genomic fidelity and screened against 1,813 clinically relevant small molecules in a high-throughput 384-well format. Drug sensitivities were quantified at the compound and drug-family levels and integrated with histologic grade, pathway-level mutational profiles, and available clinical treatment information.

**Results:** PDOs preserved hallmark tumor features, including glandular organization and subclonal mutational architecture. Pharmacotyping revealed both shared and subtype-specific vulnerabilities. Classical (well/moderately differentiated) PDOs showed enriched mutations in DNA repair, mitotic spindle, and chromatin-regulatory pathways and were preferentially sensitive to topoisomerase inhibitors, microtubule poisons, and HDAC inhibitors. In contrast, basal (poorly differentiated) PDOs displayed coordinated defects in mitochondrial function, vesicle trafficking, and ubiquitin-mediated proteostasis, at the pathway level, that conferred a previously unrecognized vulnerability to cardiac glycosides. Sensitivities to standard PDAC agents were heterogeneous across models, underscoring the limited predictive value of genotype alone and the need for functional drug testing.

**Conclusions:** This integrated genomic and pharmacologic analysis demonstrates that PDO pharmacotyping identifies biologically grounded, actionable vulnerabilities in PDAC, including novel therapeutic opportunities in basal, chemo-resistant tumors. These findings support PDO-guided functional profiling as a clinically relevant platform for refining drug selection and expanding treatment options for patients with PDAC.

**Significance:** PDAC is dominated by chemoresistance and lacks reliable genomic predictors of therapy response. By integrating high-throughput drug screening with mutation-informed pathway analysis in patient-derived organoids, we identify differentiation-linked therapeutic liabilities, including a previously unrecognized vulnerability to cardiac glycosides in basal PDAC. These results highlight PDO pharmacotyping as a powerful functional complement to genomics for guiding treatment selection in pancreatic cancer.

## Introduction

Pancreatic ductal adenocarcinoma (PDAC) is a malignancy with one of the poorest prognoses among solid tumors. Epidemiological data indicate that PDAC is currently the seventh leading cause of cancer-related mortality worldwide [1], and projections suggest it will become the second leading cause of cancer death in developed countries within the next decade [2]. The median survival for advanced PDAC remains less than one year, and curative resection is feasible in only 10–15% of patients at diagnosis. Despite substantial research investment, there has been little improvement in outcomes compared to other solid tumors.

Comprehensive sequencing efforts have revealed the genomic hallmarks of PDAC, including near-universal KRAS mutations and frequent alterations in TP53, CDKN2A, and SMAD4 [3]. However, unlike cancers driven by targetable kinases or hormone receptors, these canonical PDAC mutations remain largely non-actionable with current therapies. Only a small subset of patients with specific KRAS variants, such as KRAS G12C, may be eligible for emerging targeted agents that have shown early clinical activity [4]. As a result, genomics alone has provided limited benefit for most patients, underscoring the need for functional models that move beyond descriptive sequencing to identify therapeutically relevant vulnerabilities.

One promising approach is drug repurposing, which involves systematically testing clinically approved agents for other indications in alternative oncologic contexts. Since these agents have established safety and pharmacokinetic profiles, repurposing can accelerate translation to the clinic, bypassing years of early-phase development. Historically, repurposed drugs have had a transformative impact, as seen with thalidomide in multiple myeloma [5], arsenic trioxide, and retinoic acid in acute promyelocytic leukemia [6, 7]. More recently, interest has grown in evaluating widely used agents such as metformin and aspirin as anticancer therapies [8–10], as well as statins, which have been associated with reduced cancer incidence and mortality in observational studies [11–13]. Using this strategy, we identified digoxin as a promising and effective therapy for a rare and aggressive type of endometrial cancer [14] and fludarabine phosphate as a potential therapeutic for N-MYC-overexpressing neuroendocrine prostate cancer [15]. These studies highlight how drug repurposing, when paired with functional modeling, can uncover unexpected and clinically actionable opportunities.

In PDAC, where effective therapies remain scarce, patient-derived models provide a unique opportunity to apply this paradigm. Traditional experimental models have specific limitations: established cell lines lack sufficient heterogeneity and undergo culture adaptation [16], while patient-derived xenografts (PDXs) are costly, time-consuming, and unsuitable for high-throughput pharmacologic interrogation [17]. In contrast, patient-derived organoids (PDOs) lack these critical deficiencies. PDOs retain the histologic, molecular, and clonal features of the matching parental tumors [18, 19], can be expanded and banked, and are compatible with high-throughput drug screening. In PDAC, PDOs have already demonstrated the ability to predict patient response to chemotherapy [20], positioning them as functional avatars for translational oncology. Here, we utilize PDOs to conduct large-scale drug-repurposing screens of over 1,800 FDA-approved compounds, revealing both patient-specific vulnerabilities and generalizable therapeutic opportunities. By combining PDO fidelity with the power of pharmacotyping, we establish a platform that directly addresses the limitations of genomics as a stand-alone approach, fast-tracks therapeutic discovery, and lays the groundwork for PDO-guided clinical translation in PDAC.

## Results

### Establishment of a PDO Collection Capturing the Morphologic Spectrum and Epithelial Polarity of Pancreatic Ductal Adenocarcinoma

We successfully established 10 independent PDO models from surgically resected PDAC specimens, capturing the spectrum of tumor differentiation observed clinically. Overall, these 10 organoids were derived from 14 tumor samples, yielding a success rate of ∼70%, consistent with prior reports of PDO derivation in gastrointestinal malignancies [21–23]. Tumors were enzymatically dissociated, and the resulting cells were embedded in Cultrex Basement Membrane Extract (BME) and cultured in serum-free media optimized for PDAC organoid expansion [24]. Organoids typically formed within 5-7 days and were successfully expanded over multiple passages. To assess morphologic fidelity, we performed direct histologic comparisons of each PDO model and its matched parental tumor using hematoxylin and eosin (H&E) staining. Tumor-PDO pairs were ranked by clinical histologic grade (well, moderate, or poor differentiation; **Figure 1**) based on systematic evaluation of glandular architecture, epithelial polarity, and nuclear features.

**Figure 1.**
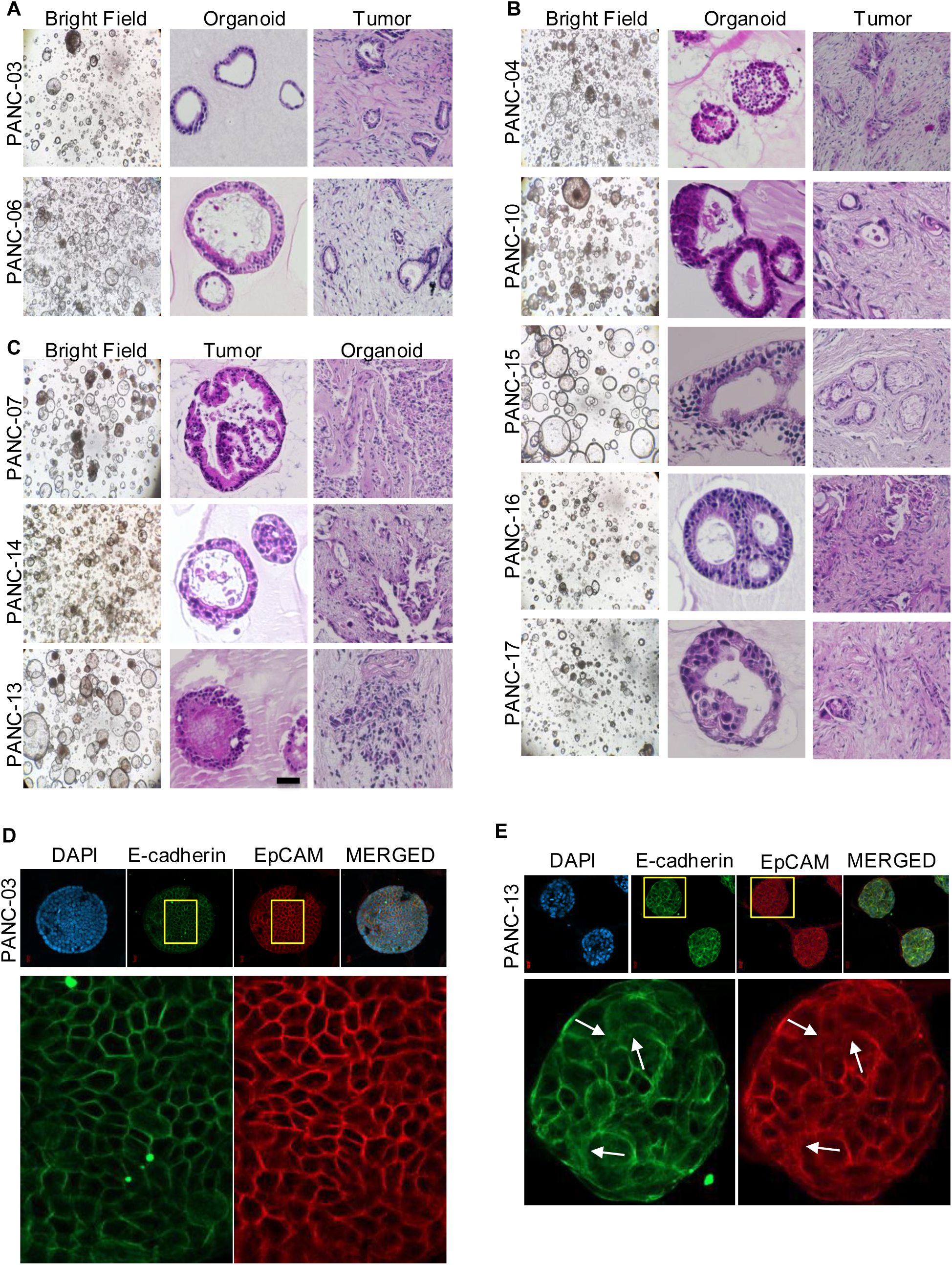
Representative images of patient-derived PDAC organoids and matched primary tumors. (A, B, and C) For each PDO model, the left panels show bright-field micrographs of 3D organoid cultures (“Bright Field”), the middle panels show histological images of the corresponding patient tumor section (“Tumor”), and the right panels show histology of the PDO model (“Organoid”). (A) Well-differentiated PDAC samples. (B) Moderately differentiated PDAC samples. (C) Poorly differentiated PDAC samples. (D) Representative fluorescence images of the PANC-03 organoid model stained with fluorescently labeled antibodies for cell-cell adhesion protein E-cadherin (green) and epithelial cell marker EpCAM (red), with DAPI nuclear counterstain. (E) The same analysis applies to the PANC-13 PDO line. The arrows indicate cells displaying scattered cytoplasmic or patchy membrane staining.

PDOs derived from Grade 1 tumors exhibited classic ductal gland morphology with strong architectural fidelity to their tumors of origin. Samples from both PANC-03 and PANC-06 tumors formed large, cystic organoids composed of a single epithelial layer surrounding a central lumen. The tumor tissues showed well-formed glands with clear apical-basal polarity, which was recapitulated in the PDOs: nuclei remained basally located, with low pleomorphism and cytoplasmic clarity **(Figure 1A)**. These PDOs displayed minimal mitotic activity, as assessed by nuclear staining, and retained the hallmark features of well-differentiated PDAC [20].

PDOs from Grade 2 tumors exhibited a diverse range of intermediate morphologies. PANC-04, PANC-10, and PANC-15 formed organoids with partially defined lumina and areas of multilayered epithelial stratification. In PANC-16 and PANC-17, gland-like regions were present but interspersed with solid or compact areas that lacked clear polarity. These features mirrored their matching tumor sections, which showed disrupted ductal organization and increased nuclear irregularity **(Figure 1B)**. Notably, PANC-07, representing an IPMN-associated carcinoma, displayed a mixed papillary-glandular architecture consistent with the clinical diagnosis. The matching PDO showed partial luminal formation, reflecting the tumor’s complex differentiation state **(Figure 1C)**. Across this group, PDOs captured the architectural variability and moderate cytologic atypia characteristic of G2 tumors.

The PANC-14 tumor was annotated as moderately to poorly differentiated, and its corresponding PDO displayed a hybrid morphology. Organoids exhibited partial lumen formation in some areas and dense, solid aggregates in others. Epithelial polarity was inconsistently maintained, and nuclear features were more pleomorphic compared to G2 PDOs. This intermediate phenotype matched the mixed histology observed in the tumor, supporting the capacity of PDOs to recapitulate transitional states between classic and basal-like PDAC [20]. The PDO derived from the Grade 3, PANC-13 tumor, propagated in densely packed, solid spheroids with no visible lumen and complete loss of epithelial polarity **(Figure 1C)**. The tumor section showed sheets of undifferentiated cells with marked nuclear pleomorphism and the absence of gland formation, features faithfully reflected in the corresponding PDO. These morphologic traits define the poorly differentiated PDAC phenotype and demonstrate that even high-grade architecture is preserved in culture [24].

Taken together, these side-by-side comparisons confirm that our PDO collection recapitulates the full morphologic continuum of PDAC, from well-formed ductal glands to disorganized solid masses. Organoid structures reflected the degree of differentiation, polarity, and cytologic identity of their tumors with striking consistency **(Table 1)**. These findings align with prior studies showing that PDAC organoids preserve the glandular architecture and epithelial lineage markers of the original tumors [25]. This level of morphologic concordance underscores the utility of PDOs as physiologically faithful models for downstream molecular and pharmacologic profiling.

**Table 1:** Summary table of clinicopathologic and histologic features. . The table lists each PDO model (PANC-03, PANC-06, etc.), the matching expert clinical diagnosis and tumor grade, the tumor’s histologic subtype (e.g. Classical, Basal, Hybrid), and the PDO’s morphological classification. A “Yes” indicates concordance between tumor and reference (clinical or organoid) for each category. All PDOs retain the differentiated subtype of the original tumor, and tumor/PDO classifications match (as indicated in the final column). NOS- not otherwise specified.

| PDO ID | Expert Clinical Diagnosis | Tumor Morphology | PDO Morphology | Tumor vs Clinical | PDO vs Clinical | Tumor vs PDO |
| --- | --- | --- | --- | --- | --- | --- |
| PANC-03 | G1, Well differentiated Ductal adenocarcinoma (NOS) | Well differentiated | Well differentiated | Yes | Yes | Yes |
| PANC-06 | G1, Well differentiated Ductal adenocarcinoma (NOS) | Well differentiated | Well differentiated | Yes | Yes | Yes |
| PANC-04 | G2, Moderately differentiated Ductal adenocarcinoma (NOS) | Moderately differentiated | Moderately differentiated | Yes | Yes | Yes |
| PANC-10 | G2, Moderately differentiated Ductal adenocarcinoma (NOS) | Moderately differentiated | Moderately differentiated | Yes | Yes | Yes |
| PANC-15 | G2, Moderately differentiated Ductal adenocarcinoma (NOS) | Moderately differentiated | Moderately differentiated | Yes | Yes | Yes |
| PANC-16 | G2, Moderately differentiated Ductal adenocarcinoma (NOS) | Moderately differentiated | Moderately differentiated | Yes | Yes | Yes |
| PANC-17 | G2, Moderately differentiated Ductal adenocarcinoma (NOS) | Moderately differentiated | Moderately differentiated | Yes | Yes | Yes |
| PANC-07 | G2, Moderately differentiated, (IPMN w/ invasive carcinoma) | Moderately differentiated | Moderately differentiated | Yes | Yes | Yes |
| PANC-14 | Moderate to poorly differentiated Ductal adenocarcinoma (NOS) | Poorly differentiated | Poorly differentiated | Yes | Yes | Yes |
| PANC-13 | G3, Poorly differentiated Ductal adenocarcinoma | Poorly differentiated | Poorly differentiated | Yes | Yes | Yes |

To corroborate the morphologic and phenotypic classifications derived from histopathology, we evaluated the epithelial integrity and polarity of the PDO collection using immunofluorescence staining for E-cadherin and EpCAM, canonical junctional and epithelial markers of pancreatic ductal differentiation, and nuclear counterstaining with DAPI. Across the collection, all PDOs exhibited membrane-associated expression of both E-cadherin and EpCAM, confirming maintenance of epithelial identity and exclusion of mesenchymal drift during culture. Specifically, immunofluorescence of the well-differentiated PANC-03 organoids revealed strong, co-localized E-cadherin and EpCAM staining along the lateral borders of contiguous cells **(Figure 1D)**. Both markers formed a continuous ring around the cell membranes, indicating tight cell-cell contacts. Indeed, E-cadherin’s role in maintaining epithelial junctions and polarity is well established [26, 27], and EpCAM forms homophilic adhesions that preserve epithelial integrity [26]. The sharply defined junctional localization of both proteins in PANC-03 confirms that these cells retain a fully epithelial phenotype.

In contrast, the poorly differentiated PANC-13 organoids displayed markedly diffused staining in the junctions, compared to PANC-03. E-cadherin and EpCAM signals were more discontinuous in PANC-13, with many cells displaying only scattered cytoplasmic or patchy membrane staining **(Figure 1E)**. Rather than forming smooth rings at cell borders, both markers appeared focal and uneven. This loss of uniform junctional staining reflects a breakdown of epithelial polarity, consistent with partial epithelial-to-mesenchymal transition. Such disorganization of E-cadherin and EpCAM is documented in high-grade PDAC, with multiple studies reporting reduced or mislocalized membranous E-cadherin in poorly differentiated, non-cohesive tumors [26, 27]. Likewise, loss of EpCAM accompanies dedifferentiation in carcinomas [28]. Thus, the patchy cytoplasmic EpCAM and attenuated E-cadherin in PANC-13 reflect its origin from an undifferentiated tumor, matching the parental tumor grades.

### PDOs Recapitulate Genomic Architecture and Subclonal Complexity of Parental Tumors

To rigorously assess the genomic fidelity of our PDOs, we performed whole-exome sequencing (WES) on paired tumor and organoid samples and conducted a multi-layered comparative analysis. The first step involved profiling somatic variants in each tumor-PDO pair to determine the extent of shared mutations. In nearly all cases, a high proportion of mutations detected in the tumor were also present in the corresponding PDO, with an average concordance of approximately 80–90% of single-nucleotide variants (SNVs) **(Figure 2A)**. This is consistent with prior study demonstrating that PDOs retain most clonal somatic mutations from their parental tumors [29].

**Figure 2.**
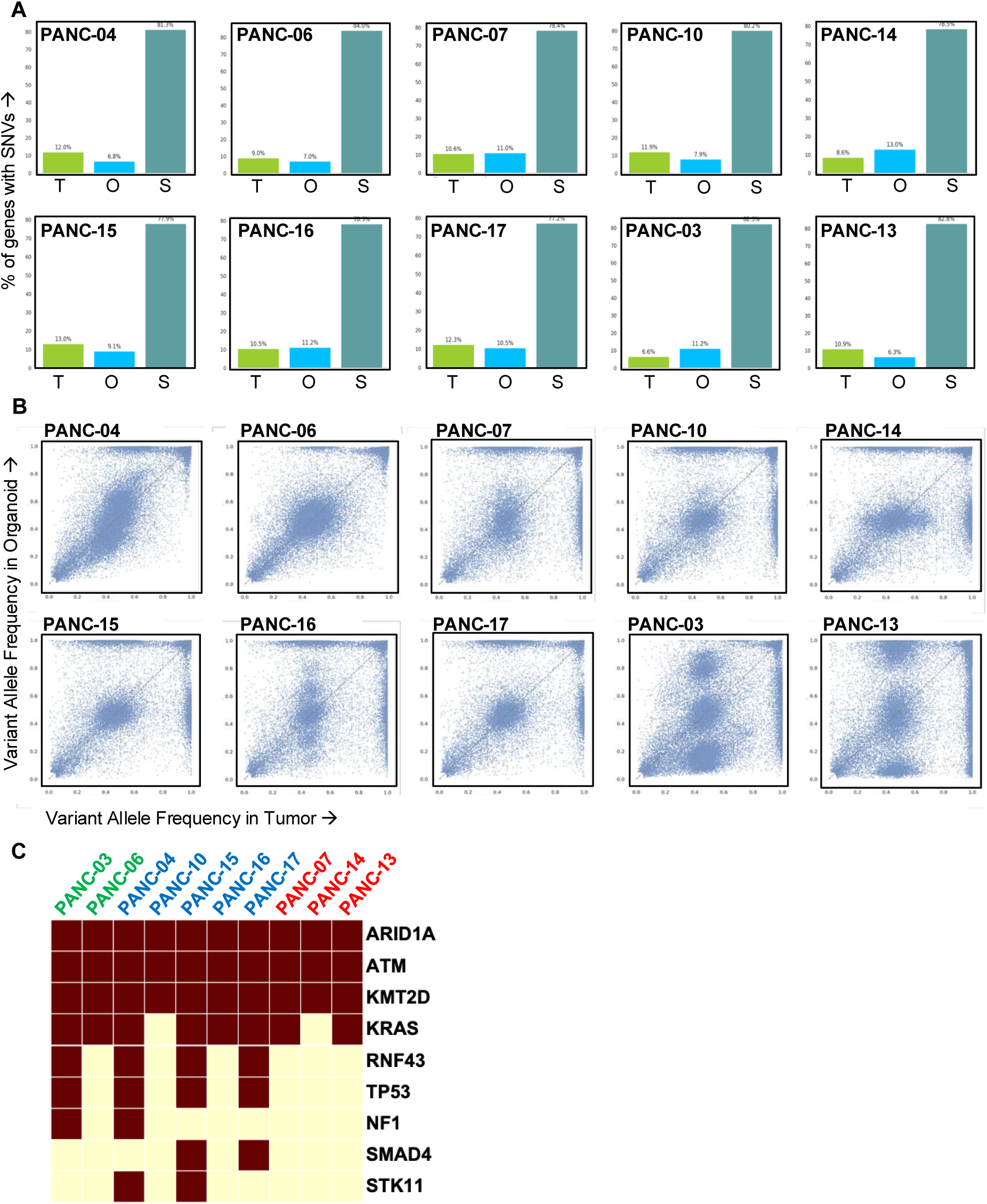
Somatic variant overlap between tumors and PDOs. (A) Bar plots showing, for each individual PDAC case, the percentage of mutated genes found exclusively in the tumor (T), exclusively in the matching PDO model (O), or in both (shared, S). (B) Scatter plots of variant allele frequencies for each somatic mutation detected in an individual PDAC tumor versus matching mutations in the matched PDO model. Each point represents one variant; the close correlation (points near the diagonal) indicates that PDOs retain the mutational spectrum of the original tumors. (C) Heatmap displaying the mutational landscape of core PDAC driver genes across the PDO cohort. The conserved driver mutation signatures reinforce the genomic fidelity of PDOs and support their use as functional avatars for pharmacologic interrogation.

We next examined whether the mutational processes shaping the tumor genome were recapitulated in organoid culture **(Supplementary Table 1)**. Mutation spectra stratified by base-substitution class revealed highly similar trinucleotide profiles between tumors and matched PDOs. Specifically, C>T transitions were the predominant SNV class in both tumors and PDOs, reflecting the age-related deamination signature that characterizes PDAC. Other substitution classes, including C>A, T>C, and T>G changes, were also preserved at similar proportions **(Supplementary Figure 1)**. These results suggest that the biological mutational processes operative in the primary tumors are most likely retained in the derived organoid cultures, further supporting molecular fidelity at the level of mutational mechanism.

To investigate subclonal fidelity, we compared variant allele frequencies (VAFs) between tumors and matched PDOs for all shared SNVs **(Figure 2B)**. In most cases, including PANC-04, PANC-06, PANC-07, PANC-10, PANC-14, PANC-15, PANC-16, and PANC-17 VAF scatter plots showed a tight diagonal correlation, indicating preservation of both dominant and minor subclonal populations. As expected, VAFs were modestly higher in PDOs due to the absence of stromal architecture and increased tumor cell purity in culture, a phenomenon reported previously in organoid sequencing studies [30, 31]. These data demonstrate that PDOs do not simply retain a subset of clones but instead preserve the full subclonal spectrum of the tumor, thereby improving sensitivity for detecting low-frequency variants. Two cases (PANC-03 and PANC-13) showed deviations from this trend, with VAF distributions that diverged from the tumor diagonal **(Figure 2B)**. In both cases, all tumor-derived variants were retained in the PDOs, but their relative VAFs shifted slightly compared with those in the parental tumors. Thus, these shifts in clonal frequency reflected enhanced representation rather than the absence of tumor-derived variants.

Finally, we examined the distribution of canonical PDAC driver mutations across the PDO cohort. High-allele-frequency variants were consistently preserved, with recurrent alterations detected in ARID1A (10/10 PDOs), ATM (10/10 PDOs), and KMT2D (10/10 PDOs), alongside activating KRAS mutations (8/10 PDOs) and inactivating mutations in TP53 (4/10 PDOs), RNF43 (4/10 PDOs) and SMAD4 (2/10 PDOs). Notably, CDKN2A, a classical PDAC tumor-suppressor, was not mutated in our cohort, suggesting biological heterogeneity in the patient tumors rather than a limitation of the PDO system. This pattern indicates that major trunk driver events, those acquired early during pancreatic tumor evolution, are faithfully maintained during PDO derivation and expansion **(Figure 2C)**. In conclusion, the WES-based comparison of tumor-PDO pairs demonstrates that PDOs preserve the full genetic blueprint of their parental PDAC tumors. Both mutation content and base-substitution spectra were maintained, and subclonal populations were conserved without evidence of clonal loss. These findings confirm that PDOs are robust, genomically faithful models of PDAC suitable for functional studies, molecular stratification, and drug-response prediction.

### Scalable Pharmacologic Screening in Pancreatic PDOs Identifies Distinct Drug-Response Signatures

Having established the morphologic and genomic fidelity of our pancreatic PDO collection, we next leveraged this platform for functional pharmacologic interrogation. Each organoid model derived from a patient tumor is challenged with a library of pharmacologic agents and scored for sensitivity based on viability loss **(Figure 3A)**. This approach recapitulates precision oncology principles at the *ex vivo* level, enabling patient-specific prediction of therapeutic vulnerabilities.

**Figure 3.**
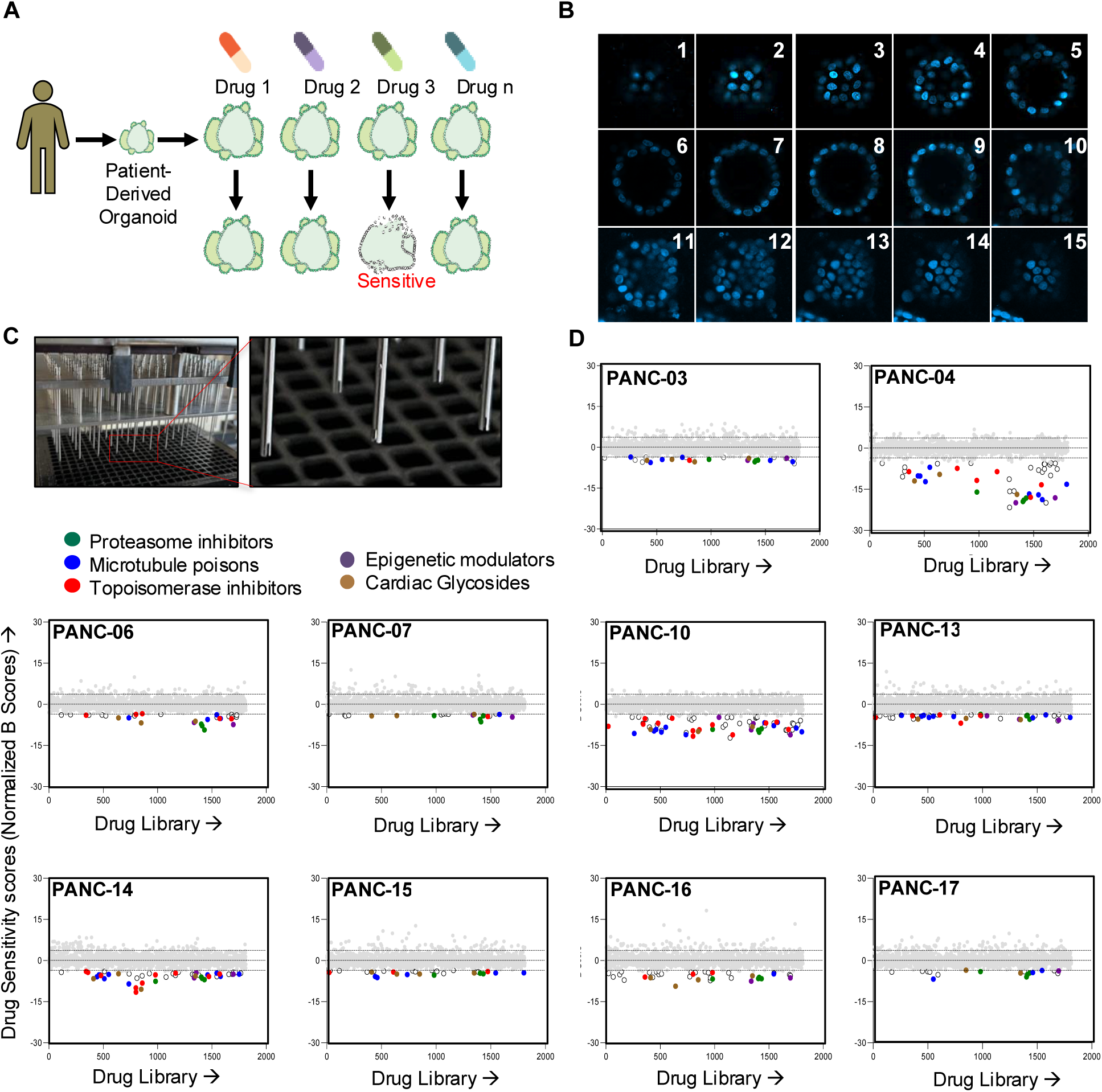
High-throughput pharmacologic profiling of PDAC PDOs identifies drug classes with selective sensitivity. (A) Schematic overview of the organoid-based drug screening strategy. Patient-derived organoids (PDOs) were exposed to a library of 1,813 clinically approved small molecules. Viability was measured after 7 days of treatment to identify compounds that preferentially suppressed growth of individual organoid models. (B) Confocal imaging of PDO spheroid formation in 384-well format. Representative confocal DAPI-stained z-stack images (5-µm intervals) demonstrate that pancreatic PDOs form uniform, lumen-containing epithelial spheres under miniaturized screening conditions. Nuclear organization and spheroid morphology remain intact across the full z-axis, confirming that PDO structural integrity is preserved during high-throughput assay plating. Numbers represent the sequence of z-axis images. (C) Images show the stainless-steel pin-tool array used for nanoliter-scale compound transfer into 384-well PDO plates. The system deposits 200 nL of each drug with high positional accuracy, enabling reproducible and cost-efficient screening of 1,813 compounds in parallel. Insets illustrate the pin geometry and contact-based delivery mechanism. (D) Drug sensitivity profiles for each PDO model based on normalized B-scores. Each dot represents a single compound; the major drug families among the top hits are highlighted: proteasome inhibitors (green), microtubule poisons (blue), topoisomerase inhibitors (red), epigenetic modulators (purple), and cardiac glycosides (brown). Compounds with more negative B-scores exhibit higher cytotoxicity than those with a full drug distribution. The two dotted horizontal lines denote statistical thresholds used to identify outlier compounds. Points below these cutoffs represent drugs with responses that exceed expected assay variability and are considered significant hits.

To ensure that miniaturized high-throughput assays faithfully recapitulated 3D organoid architecture, we first verified spheroid formation directly in the 384-well screening plates. Confocal DAPI z-stack imaging at 5-µm intervals **(Figure 3B)** demonstrated that all PDO models reproducibly formed uniform, well-organized spherical structures with intact nuclear organization. These data confirm that the organoids maintain physiologic 3D architecture under screening conditions, providing a robust foundation for downstream functional assays. To further support assay scalability and precision, we employed a pinning tool that dispenses ∼200 nL of compound per well with high positional accuracy **(Figure 3C).** This system enables parallel delivery of 1,813 compounds into 384-well plates with minimal volume error, ensuring uniform drug exposure across PDOs. Together, the imaging and dispensing workflows establish a reliable, quality-controlled platform for large-scale pharmacologic screening. Each of the ten PDO models was screened against the Targetmol Bioactive Compound Library, comprising 1,813 small molecules that span a wide range of therapeutic mechanisms [32]. Of these compounds, approximately 61% are FDA-approved drugs, while 38% are approved by regulatory agencies in other countries. Notably, the library also includes nearly 50 compounds currently in clinical trials, enhancing its relevance for translational investigations. To our knowledge, this represents the first application of the full Targetmol library in PDAC PDOs, providing an unbiased, broad pharmacologic landscape of pancreatic tumor vulnerabilities. PDOs were plated in the 384-well format and exposed to each drug at a single concentration of 2 µM for seven days. Viability of PDO cells was assessed using the luminescent 3D CellTiter-Glo assay, which measures intracellular ATP as a proxy for metabolic activity. Although single-dose screening lacks detailed dose**-**response curve resolution, this approach offers a cost-efficient and scalable strategy for identifying high-confidence hits and prioritizing compound classes for more detailed follow-up studies. Given the library’s scope (>1,800 compounds plus DMSO controls) and the limited throughput of 3D organoid assays, single-dose profiling provides a scalable approach to identify Grade-dependent and tumor-specific sensitivities.

Quantitative drug sensitivities, normalized as B-scores, are shown for each PDO model **(Figure 3D)**. While most compounds fell within the neutral response range, each PDO model exhibited selective vulnerabilities to specific drug classes. Despite being exploratory, the single-dose screen captured biologically meaningful differences across individual PDO models. Several models exhibited low overall sensitivity, consistent with a chemoresistant phenotype, while others (e.g., PANC-04 or PANC-10) showed broader responses across compound classes **(Figure 3D)**. These data support the hypothesis that PDAC subtypes have distinct pharmacologic liabilities that can be uncovered through high-throughput organoid screening.

### Grade- and Subtype- Linked Pharmacologic Vulnerabilities in PDAC PDOs

To characterize pharmacologic vulnerabilities, we classified compounds by mechanisms of action and quantified the frequency with which each class was represented among the top-performing agents using a weighted enrichment metric. Across all ten PDO models, proteasome inhibitors and epigenetic modulators displayed the highest overall enrichment among top cytotoxic responses, followed by microtubule poisons, cardiac glycosides, and topoisomerase inhibitors **(Figure 4A)**. Antimetabolites and general cytotoxic drugs were rarely represented among the strongest responses. Radar plot analysis **(Figure 4B)** further demonstrated that proteasome and HDAC inhibitors elicited broad sensitivity across models, whereas microtubule poisons and topoisomerase inhibitors showed greater heterogeneity. Examination of the five most potent compounds per PDO model reinforced these trends, with recurrent representation of proteasome inhibitors (carfilzomib, MLN-2238, MLN-9708), HDAC inhibitors (romidepsin, panobinostat), microtubule poisons (vincristine), and topoisomerase inhibitors (10-hydroxycamptothecin, topotecan, daunorubicin) **(Supplementary Fig. 2)**.

**Figure 4.**
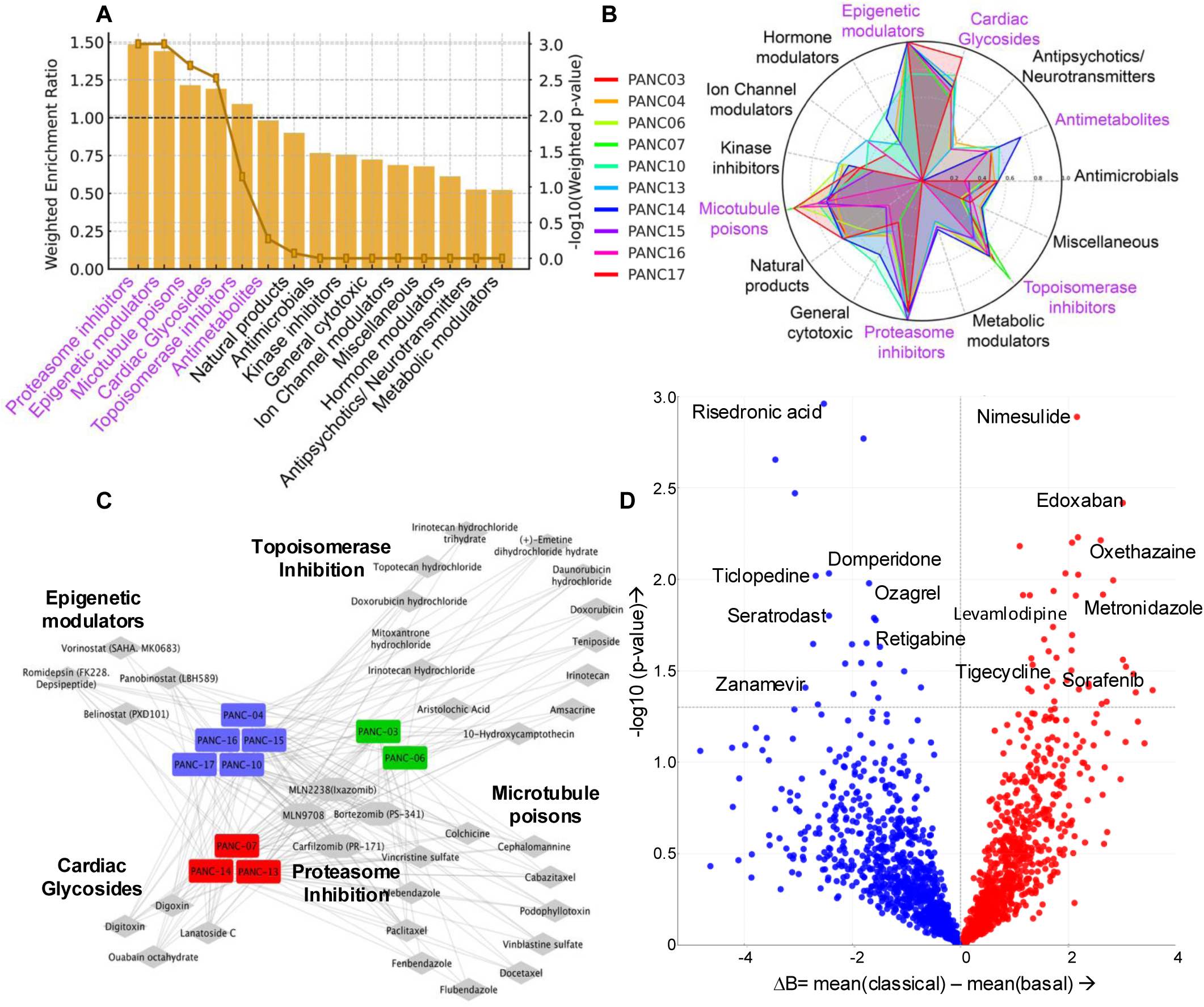
Network analysis of PDO-specific drug hits reveals differentiation-stratified pharmacologic vulnerabilities. (A) Enrichment analysis of drugs’ sensitivities by pharmacologic class combined for all tested PDO models. Drug families were ranked by weighted enrichment ratio (bars, left y-axis), with statistical significance indicated by the −log10(p-value) (line, right y-axis). Proteasome inhibitors, epigenetic modulators, microtubule poisons, cardiac glycosides, and topoisomerase inhibitors emerged as the most enriched classes across the PDO cohort. The dashed horizontal line denotes the neutral enrichment threshold (WER = 1.0), corresponding to the expected frequency of hits if compounds were randomly distributed across drug families. Drug classes exceeding this threshold are considered significantly enriched for high-activity compounds, indicating preferential sensitivity in the PDO cohort. Classes falling below the line show no enrichment. (B) Radar plot summarizing mean sensitivity across drug families for all ten PDO models. Each line represents one model, colored by model ID. Axes correspond to drug-family categories, and radial distance from the center reflects the normalized mean B-score for all compounds within that family, where higher values indicate greater sensitivity and values near the center indicate lower or minimal activity. Classes with the largest outward displacement, such as proteasome inhibitors, epigenetic modulators, and cardiac glycosides, represent drug families with the strongest shared activity across PDOs. In contrast, inward-pointing values reflect variable or limited sensitivity. (C) A drug–PDO interaction network generated using Cytoscape connecting each of the ten pancreatic ductal adenocarcinoma (PDAC) organoid models to their top five most active compounds, grouped by pharmacologic classes. Nodes are colored based on organoid differentiation state: well-differentiated (green), moderately differentiated (blue), and poorly differentiated (red). Drug nodes are labeled and clustered by functional class: proteasome inhibitors, epigenetic modulators, topoisomerase inhibitors, microtubule poisons, and cardiac glycosides. This visualization demonstrates that proteasome inhibitors exhibit broad activity across all PDOs, whereas topoisomerase inhibitors and microtubule poisons are more strongly associated with well-differentiated models (PANC-03, PANC-06). Moderately differentiated PDOs (e.g., PANC-04, PANC-15, PANC-17) are enriched for epigenetic modulators, while poorly differentiated PDOs (PANC-13, PANC-14) uniquely associate with cardiac glycosides. These patterns illustrate how histologic differentiation correlates with specific drug class sensitivities. (D) Volcano plot illustrating differential drug responses between classical (n=7) and basal (n=3) PDOs. For each compound, the x-axis shows the difference in mean B-scores (ΔB = mean(classical) – mean(basal), where negative values (blue) indicate preferential cytotoxicity in classical PDOs and positive values (red) indicate stronger cytotoxicity in basal PDOs. The y-axis represents –log10(p-value) from Welch’s t-test. Dashed lines denote ΔB = 0 and p = 0.05. This representation captures the broad asymmetry in response patterns and visually separates compounds with preferential activity toward classical versus basal PDO models.

To determine whether histological differentiation influences drug sensitivity, we applied a rank-based enrichment framework to identify mechanistic classes (drugs with similar mechanisms of action) consistently represented among the most compounds in each PDO model. For each model, compounds were ordered by increasing B-score, the top 5, 10, 15, and 20 strongest cytotoxic responses were extracted, and enrichment was calculated as the frequency with which members of each mechanistic class appeared within these top-ranked sets. Enrichment values were then averaged across well-, moderately-, and poorly differentiated PDO types to identify grade-associated patterns.

Although proteasome inhibitors remained broadly enriched across all differentiation types, differentiation-linked patterns emerged **(Figure 4C)**. Topoisomerase inhibitors showed greater enrichment in well-differentiated PDOs, whereas HDAC inhibitors appeared more frequently among top responses in moderately differentiated PDOs. Cardiac glycosides, though potent across the cohort, were most consistently enriched among the strongest responses in poorly differentiated PDOs. Statistical testing using Kruskal–Wallis analyses preserved these directional patterns but did not yield significant group-level differences, due to limited class sizes.

Since histologic differentiation in PDAC strongly correlates with transcriptional state, well and moderately differentiated tumors typically correspond to the classical subtype, and poorly differentiated tumors align with basal features, we next asked whether pharmacologic selectivity could be independently resolved by transcriptional subtype alone. In contrast to the rank-based approach, which identifies the most potent compounds within each PDO model, this analysis examined whether classical and basal PDOs differ in their average sensitivity to each drug across the entire library. PDO models were classified as classical (PANC03, PANC04, PANC06, PANC10, PANC15, PANC16, PANC17) or basal (PANC07, PANC13, PANC14). For each compound, we calculated the difference in mean B-score between classical and basal PDOs and performed Welch’s t-tests to identify subtype-selective responses **(Figure 4D)**. This analysis identified 35 basal-selective compounds (p < 0.05, ΔB > 0, and mean basal B-score < 0). These drugs included ROS-inducing kinase inhibitors (sorafenib, nintedanib), antimicrobials with mitochondrial or membrane-disruptive activity (metronidazole, tigecycline), membrane- and ion-channel–active agents (clofazimine, lindane, oxethazaine), and lipid/sterol-perturbing compounds such as lithocholic acid. Additional basal-selective agents, including edoxaban (suppresses PAR-mediated Ca²⁺ signaling), nimesulide (COX-2 inhibitor that induces mitochondrial/oxidative stress), levamlodipine (L-type Ca²⁺-channel blocker that disrupts calcium influx and membrane excitability), and theobromine (methylxanthine that modulates cyclic-nucleotide and Ca²⁺-linked membrane excitability), further reinforced this pattern of stress-adapted vulnerabilities. Collectively, these compounds converge on pathways involving mitochondrial stress, oxidative imbalance, membrane disruption, and perturbation of ion homeostasis, features characteristic of basal PDAC biology.

Conversely, 21 compounds were significantly classical-selective (p < 0.05, ΔB < 0, and mean classical B-score < 0). Classical PDOs were disproportionately sensitive to inhibitors of defined metabolic or receptor-mediated signaling nodes. These included neuraminidase and thromboxane-pathway inhibitors (zanamivir, ozagrel, seratrodast), COX-2 and FPPS-targeting agents (valdecoxib, risedronic acid, ibandronic acid), COMT inhibition (tolcapone), glucose-transport inhibition (dapagliflozin), and neuronal-like signaling modulators (suvorexant, dextromethorphan, retigabine). The classical-selective set was largely distinct from the top-ranked agents identified through the grade-based enrichment analysis. This difference reflects the orthogonal nature of the two approaches: the rank-based method determines the most potent drugs within each PDO model, whereas the subtype-stratified analysis captures drugs whose mean activity differs across transcriptional states. Importantly, pooling well- and moderately differentiated PDOs into the classical subtype necessarily averages their distinct sensitivities, such as topoisomerase inhibitors being preferentially active in well-differentiated models and HDAC inhibitors in moderately differentiated models, thereby diluting class-specific signals in the subtype-level comparison. This limitation is expected to be mitigated in larger cohorts, where increased sample size will permit separate stratification of well versus moderately differentiated classical tumors and more robust detection of overlapping and grade-specific drug-class enrichments.

Interestingly, across both frameworks, poorly differentiated, basal PDOs consistently demonstrated vulnerability to agents that disrupt mitochondrial function, membrane stability, redox balance, and Na⁺/K⁺-ATPase-linked ion transport. The convergence of these biological signatures across two analytically independent strategies, one emphasizing within-PDO model cytotoxicity ranking and the other contrasting subtype-wide differences in mean drug response, reveals that PDAC differentiation and molecular subtype encode common underlying pharmacologic dependencies.

### Distinct Pathway-Level Defects Underlie Differentiation-Dependent Drug Sensitivities in PDAC PDOs

Given the grade- and subtype-specific drug sensitivity patterns, we next examined how PDO models-predicted drug sensitivities aligned with the clinical treatments and outcomes of the corresponding patients **(Table 2)**. As all PDOs were derived from treatment-naïve surgical specimens, this comparison provides contextual, but not evaluative, insight into how functional drug susceptibilities relate to the therapies routinely used in PDAC. Most patients later developed recurrent disease, and although they received standard cytotoxic regiments, PDO profiling revealed a broader spectrum of potentially actionable vulnerabilities. In some cases, PDOs demonstrated sensitivity to drug classes commonly used in PDAC treatment, whereas in others, the screens identified additional opportunities, including proteasome and HDAC inhibitors, that were not part of the administered regimens. These observations underscore how PDO pharmacotyping can complement clinical decision-making by revealing therapeutic options that extend beyond standard-of-care combinations.

**Table 2.** Clinical treatment information for patients corresponding to each PDO model. Summary of treatment-naïve status, adjuvant or first-line therapy, systemic regimens administered, and second-line therapies (if applicable) for the ten PDAC patients from whom PDOs were derived. All PDOs were established from resection specimens obtained before any systemic treatment.

| PDO | Treatment-Naive Tumor (Y/N) | Adjuvant/First Line Therapy | Clinical Systemic Therapy Administered | Second-Line Therapy (if any) |
| --- | --- | --- | --- | --- |
| PANC03 | Yes | Adjuvant therapy | Irinotecan + Oxaliplatin + Leucovorin + 5-FU (mFOLFIRINOX) | nab-Paclitaxel + Gemcitabine |
| PANC06 | Yes | Adjuvant therapy | Irinotecan + Oxaliplatin + Leucovorin + 5-FU (mFOLFIRINOX) |  |
| PANC04 | Yes | Adjuvant therapy | Capecitabine + Gemcitabine | RT + Capecitabine |
| PANC10 | Yes | None | None |  |
| PANC15 | Yes | Adjuvant therapy | Capecitabine + Gemcitabine |  |
| PANC16 | Yes (prior breast cancer therapy in 2009) | Adjuvant therapy | Irinotecan + Oxaliplatin + 5-FU (mFOLFIRINOX) |  |
| PANC17 | Yes | None | None |  |
| PANC07 | Yes | None | None |  |
| PANC14 | Yes | 1st-line metastatic | nab-Paclitaxel + Gemcitabine | Oxaliplatin + Leucovorin + 5-FU (FOLFOX) |
| PANC13 | Yes | None | None |  |

To further contextualize these findings, we examined PDO responses to commonly used PDAC agents **(Supplementary Figure 3)**. Sensitivity patterns were heterogeneous but interpretable, with measurable activity in several models to agents frequently used in adjuvant and first-line settings: gemcitabine, irinotecan, or paclitaxel. However, no standard-of-care drug demonstrated uniform activity across all PDOs, and limited sensitivity to oxaliplatin and capecitabine was observed in most models. These data highlight substantial intertumoral variability and suggest that drug response cannot be accurately inferred solely from genotype or clinical practice patterns.

To explore the biological basis for the limited concordance between PDO responses and clinical treatments, we next examined whether the mutational landscape could account for subtype- and grade-specific drug sensitivities. To this end, we examined mutations that were predominantly present in each of the well-, moderate-, and poorly differentiated PDO models. Most genetic alterations in well- and moderately differentiated PDOs overlapped, and our sample size was too small to identify any specific enrichment. Interestingly, poorly differentiated PDOs harbored several additional mutations absent in well- and moderately differentiated PDOs. Therefore, we examined mutation status by grouping PDO models into the two commonly classified PDAC subgroups, namely classical and basal.

Classical subtype PDOs harbored a concentrated burden of mutations in genes involved in DNA damage repair, replication stress response, mitotic spindle regulation, and epigenetic modulation **(Figure 5A).** Notably, several classical models carried inactivating mutations in homologous recombination and DNA repair genes, including *FANCA* and *RAD51*, as well as cell-cycle checkpoint regulators such as *WEE1*. Additional alterations in *DDB1*, *RRM1*, and the p53 regulator *RPL11* further implicate defects in genome surveillance and replication fidelity. Consistent with these genomic features, classical PDOs were more vulnerable to topoisomerase inhibitors in our drug screens **(Figure 4C)**, which induce replication stress and DNA breaks that HR-deficient cells cannot effectively resolve. Classical PDOs also carried frequent mutations in mitotic spindle and microtubule-regulatory genes, including AURKB (chromosome-segregation kinase) and the centrosomal regulator *RASSF7*. Additional alterations in microtubule structural or modifying proteins, such as *TUBA1C* and *TTLL13*, indicate underlying defects in spindle integrity and microtubule stability. These features likely sensitize classical models to microtubule poisons, consistent with their heightened response to taxanes and vinca alkaloids.

**Figure 5.**
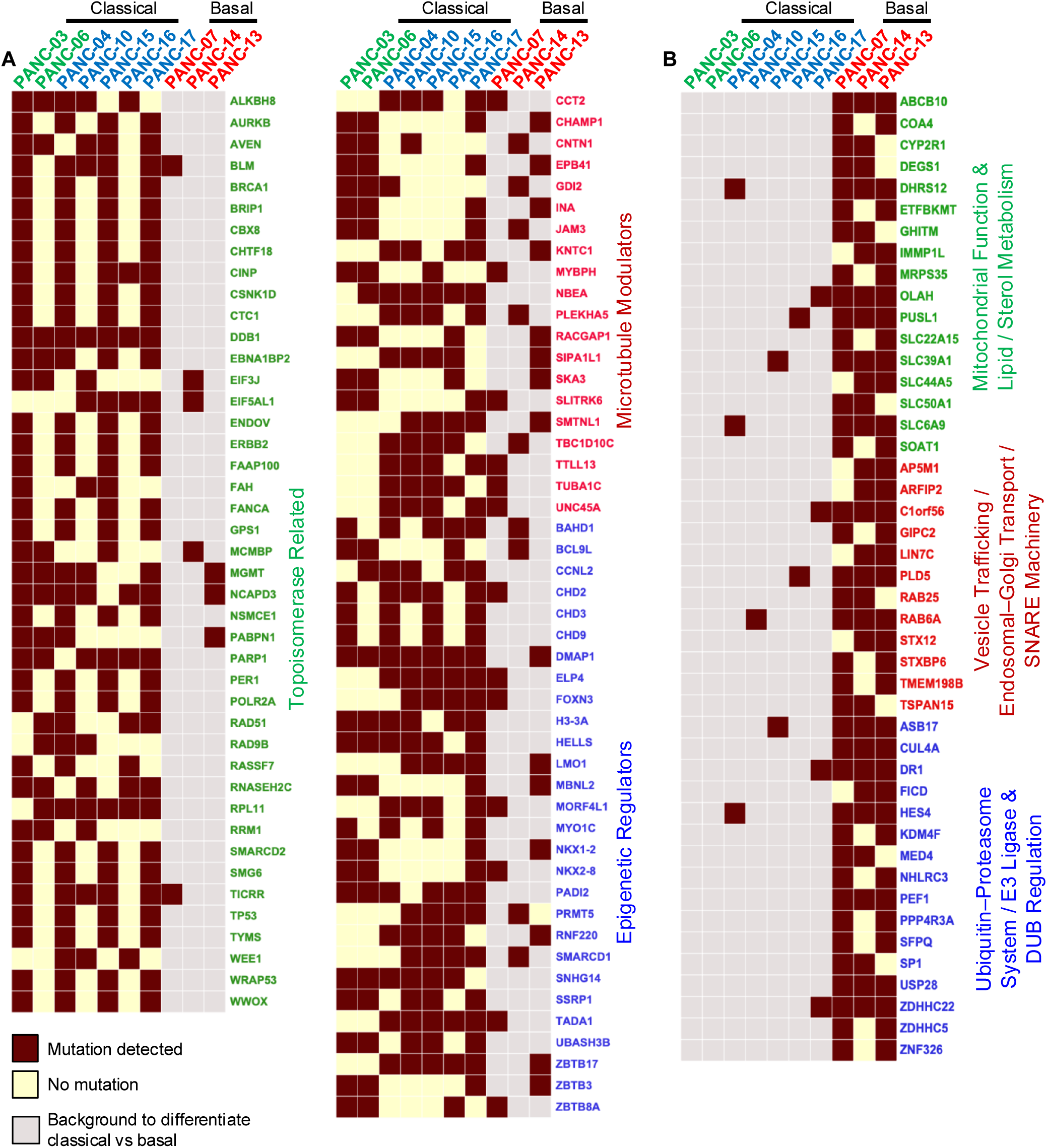
Pathway-level mutation patterns distinguishing classical and basal PDAC PDOs. (A) Heatmaps showing mutations in genes associated with DNA repair and replication stress (left), microtubule structure and mitotic regulation (upper middle), and chromatin remodeling/epigenetic control (lower middle) across ten PDO models. Columns are grouped by classical PDOs shown on the left (green/blue) and basal PDOs on the right (red). Dark squares indicate detected somatic mutations; light squares indicate wild-type status; gray shading demarcates subtype groupings. (B) Heatmaps illustrating mutations in genes involved in mitochondrial function and lipid/sterol metabolism (green), vesicle trafficking and Golgi/endo-lysosomal machinery (red), and ubiquitin–proteasome/ERAD/ligase and deubiquitinase pathways (blue).

In the epigenetic landscape, classical PDO models harbored multiple mutations in chromatin regulators, including *MORF4L1* (a component of the Tip60/NuA4 histone acetyltransferase complex), *PRMT5* (a protein arginine methyltransferase), and *SMARCD2* and *SMARCD1* (subunits of the SWI/SNF chromatin remodeling complex). These alterations suggest disrupted chromatin organization and histone modification, potentially driving an epigenetic dependency that renders these models particularly sensitive to HDAC inhibitors. This was consistent with our drug screening data, where classical PDOs (that belong to moderately differentiated grades) were also more responsive to HDAC inhibition **(Figure 4C)**. Together, these findings highlight a convergence of vulnerabilities in classical PDAC PDOs across DNA repair, mitotic control, and chromatin regulation pathways, offering a mechanistic rationale for their selective sensitivity to topoisomerase poisons, microtubule-disrupting agents, and HDAC inhibitors.

Our analyses of mutation status indicated that basal PDOs also harbor individual mutations in DNA repair pathways **(Supplementary Figure S4)**, but these did not translate into sensitivity to topoisomerase inhibitors or platinum agents. In contrast, basal PDOs displayed a distinct constellation of mutations affecting mitochondrial homeostasis, membrane lipid regulation, vesicle trafficking, and ubiquitin-mediated proteostasis pathways **(Figure 5B)**. Several basal models carried mutations in mitochondrial machinery genes such as *ABCB10* and *MRPS35*, as well as lipid metabolic regulators, including *DEGS1* and *SOAT1*. These alterations suggest impaired oxidative phosphorylation and perturbed membrane lipid composition, conditions known to exacerbate cellular stress induced by cardiac glycosides, which disrupt Na⁺/K⁺-ATPase activity and, secondarily, destabilize mitochondrial ATP production.

Importantly, basal PDOs also harbored recurrent mutations in genes linked to ubiquitin-dependent protein quality control, including components of the ER-associated degradation (ERAD) pathway and ubiquitin ligase adaptors. Mutations in several ubiquitin-pathway genes, including *CHMP1A*, *RAB25*, *RAB8A*, and *USP family* members, were enriched in basal models. These genes are central to endosome–lysosome trafficking, ubiquitin-dependent cargo sorting, and turnover of damaged proteins. Disruption of these pathways compromises the cell’s ability to clear misfolded or aggregated proteins and to regulate ion transporters via ubiquitination. This defect is directly relevant to cardiac glycoside sensitivity. Because cardiac glycosides impose strong ER and proteotoxic stress, basal PDOs, harboring mutations in ubiquitin–proteasome, ERAD, and RAB/ESCRT trafficking genes, are less capable of clearing misfolded proteins or mounting an effective stress response. Collectively, these analyses show that functional drug vulnerabilities in PDAC are shaped not simply by individual mutations but by broader pathway-level disruptions that differ markedly between classical and basal tumors. These results also underscore the power of PDO pharmacotyping to expose actionable vulnerabilities in chemo-resistant PDAC and to guide therapeutic strategies beyond standard cytotoxic regimens.

## Discussion

Our study demonstrates that PDAC PDOs faithfully recapitulate the histologic, genomic, and subclonal features of their parental tumors, which matches well the conclusions of earlier reports [33, 34]. Beyond confirming fidelity, our work shows that PDO-based functional profiling captures clinically relevant therapeutic phenotypes that genomics alone cannot predict. This is consistent with recent studies demonstrating that PDO pharmacotypes align with chemotherapy response and biomarker trajectories in treated patients [35], but our results extend this concept substantially. A central advance of this work is the scale and breadth of the screening effort. To our knowledge, this represents the largest pharmacologic repurposing screen performed in genomically validated PDAC organoids, interrogating 1,813 approved or clinically advancing agents. Unlike earlier PDO studies that primarily assessed a small panel of chemotherapies [33, 35], our unbiased high-throughput approach uncovered a landscape of unanticipated drug-class vulnerabilities, revealing that PDAC harbors targetable biochemical liabilities far beyond standard cytotoxics.

Across all models, one of the strongest and most consistent dependencies was proteasome inhibition, suggesting a general reliance on proteostasis. This is interesting because preclinical studies in PDAC models show that PS-341 (bortezomib) alone profoundly suppresses PDAC cell proliferation, and that combining bortezomib with chemotherapeutics enhances tumor suppression and extends survival [36, 37]. Similarly, several organoid models were also susceptible to HDAC inhibitors. Inhibition of HDACs or EHMT2 markedly reduces the viability of pancreatic cancer cells and remains effective in gemcitabine-resistant lines [38]. Newer inhibitors, such as CG200745, increase histone H3 acetylation, activate pro-apoptotic pathways, and sensitize resistant PDAC cells to gemcitabine therapy [39]. Although early-phase clinical trials combining proteasome inhibition and HDAC inhibition have shown negative results in unselected gemcitabine-refractory or metastatic PDAC populations [40, 41], these studies were not enriched for molecularly defined subgroups. Our results suggest that proteasome inhibitor sensitivity may not be uniform but rather highly context-dependent, reinforcing the value of functional selection for identifying patients most likely to benefit. Importantly, these vulnerabilities were not predictable from genotype alone but became clear only through functional screening.

A striking novel observation was the selective sensitivity of poorly differentiated (basal-like) PDOs to cardiac glycosides, a drug class historically unrelated to PDAC therapy. By integrating pharmacotyping with pathway-level mutation analysis, we show that basal PDAC tumors exhibit common defects in mitochondrial function, vesicle trafficking, and ubiquitin-mediated proteostasis, together creating a stress environment in which Na⁺/K⁺-ATPase inhibition, via cardiac glycosides such as digoxin, becomes cytotoxic. This represents a previously unrecognized therapeutic opportunity in the most chemoresistant PDAC subgroup. In contrast, classical (well/moderately differentiated) tumors displayed pathway-level disruptions in DNA repair, mitotic spindle regulation, and chromatin remodeling, consistent with their preferential sensitivity to topoisomerase inhibitors, microtubule poisons, and HDAC inhibitors. These differentiation-linked pharmacotypes highlight that PDAC histopathology encodes functional therapeutic vulnerabilities, a conceptual advance with direct translational implications.

The included comparison with administered clinical therapies provided contextual insight. Standard-of-care agents elicited heterogeneous responses across PDO models, mirroring the clinical reality that PDAC chemotherapy benefits only a subset of patients. Importantly, sensitivities to oxaliplatin and capecitabine were limited, underscoring the limited predictive value of genotype alone and reinforcing the need for complementary functional diagnostics.

We also acknowledge some of the key limitations of our study. The cohort size is small, the primary screen used a single-dose treatment, and PDOs lack stromal and immune components, a common disadvantage of these models. However, this was intentional to avoid culture-driven evolution and maintain maximal fidelity to the patient tumor. Thus, we designed the high-throughput screen using a single replicate at a single pharmacologically relevant concentration, expanding each PDO only to the minimal extent required for assay setup. This approach follows best practices in functional precision oncology, where excessive passaging is known to introduce transcriptional and genomic drift. By minimizing expansion, our screen captures drug sensitivities in a state that is as close as experimentally possible to the original tumor ecosystem. This unbiased screen enabled a scalable assay with the integration of pharmacologic, genomic, and differentiation-based analyses distinguishing this work from all prior PDAC PDO studies. Most importantly, our findings reveal a conceptually new direction for functional precision medicine in PDAC, indicating that tumor differentiation state and pathway-level biology, not single-gene alterations, determine drug susceptibility, enabling personalized therapeutic strategies even for chemo-refractory basal PDAC.

## Materials and Methods

### Patient Samples

Tumor samples were obtained from pancreatic cancer patients undergoing surgical resection, with the assistance of the University of Saskatchewan (USask) Biobank. All samples were pathologically confirmed. Written informed consent was acquired from all patients for the use of their clinical data and surgical specimens, as well as for the publication of potentially identifiable clinical information. This study was conducted in compliance with national guidelines and was approved by the Biomedical Research Ethics Board (Bio-REB, Approval number: 3382).

### WRN Conditioned medium

L-WRN cells (ATCC, CRL-3276) were cultured in DMEM (Cytiva, SH30243.01) with 10% FBS (Corning, MT35077CV). 0.5 mg/mL G418 (Gibco, 10131035) and 0.5 mg/mL hygromycin B (Gibco, 10687010) were added to maintain selection until cells reached confluence. The cells were split in a 1:5 ratio. G-418 and hygromycin B were removed to prevent the carryover of drugs in the conditioned medium. The cells were cultured until overconfluent and then replaced with primary culture medium (Advanced DMEM/F12 (Gibco, 12634010); 20% FBS (Corning, MT35077CV); 1% Penicillin-Streptomycin (GE Healthcare, SV30010); 2 mM L-Glutamine (Gibco, 25030081)). After 24 hours of incubation, the medium was centrifuged at 2000 g for 5 minutes to collect the supernatant. The collected conditioned medium was pooled together, filtered, aliquoted and stored in −20°C[42].

### Preparation of Complete Growth Medium (CGM)

For the expansion and maintenance of pancreatic cancer organoids, a defined Complete Growth Medium (CGM) was prepared by combining WRN-conditioned medium (25 mL) with a basal formulation of Advanced DMEM/F12 (22 mL; Gibco, Cat# 12634010) supplemented with the following additives: 1× Antibiotic-Antimycotic (Gibco, Cat# 15240062), 2 mM Glutamax (Gibco, Cat# 35050061), 10 mM HEPES (GE Hyclone, Cat# SH30237.01), and 1× B-27 supplement (Gibco, Cat# 17504044). To enhance organoid viability and support epithelial differentiation, the medium was further supplemented with 1 mM nicotinamide (Sigma, Cat# N0636-100G), 1.25 mM N-acetyl-L-cysteine (Sigma, Cat# A7250-5G), 50 ng/mL recombinant human EGF (Stemcell Technologies, Cat# 78006.1), and 100 ng/mL recombinant human FGF-10 (Peprotech, Cat# 100-26-50UG). Small-molecule inhibitors were also included to modulate key signaling pathways: 0.5 μM A83-01 (TGF-β receptor inhibitor; Selleckchem, Cat# S7692), 10 nM Gastrin (Sigma, Cat# SCP0152-1MG), and 2.5 μM CHIR99021 (GSK3β inhibitor; Sigma, Cat# SML1046-5MG). All components were sterile-filtered and stored at 4°C for short-term use or −20°C for long-term storage.

### Organoid culture

For surgical samples, the collection medium was removed, and the tissue was rinsed several times with ice-cold sterile D-PBS. The tissue was then minced into small fragments using a sterile blade. A portion of these fragments was reserved for DNA/RNA extraction or histology. The remaining fragments were transferred to a 15 mL conical tube and washed with approximately 10 mL of ice-cold sterile D-PBS. This washing step was repeated 5–10 times until the PBS supernatant appeared clear. After removing most of the final PBS wash, the tissue fragments were resuspended in 5 mL of Advanced DMEM/F12 supplemented with 500 μL of Liberase TH (500 μg/mL, Roche, 5401151001). The suspension was incubated for 1 hour at 37°C with continuous agitation and periodic pipetting to facilitate dissociation. The resulting supernatant was passed through a cell strainer with 100 μm pore size (Corning, C352360) into a 50 mL conical tube. The dissociated cells were pelleted by centrifuging at 400 × g for 5 minutes. After cell counting, the pellet was resuspended in pre-thawed, cold BME gel (R&D Systems, 3533-010-02). Then, 50 μL of the cell suspension in BME gel (containing approximately 5 × 10^5^ cells) was seeded into each well of a 37°C pre-warmed 24-well plate. The plate was inverted and incubated at 37°C for 15-20 minutes to allow the BME domes to solidify. Following this, 500 μL of 37°C pre-warmed complete growth medium was added to each well. The medium was subsequently changed every 2-3 days. Organoids were cryopreserved in freezing medium (RPMI 1640 + 20% FBS + 10% DMSO) and were routinely tested for Mycoplasma contamination.

### Histology and H&E staining

An aliquot of Histogel (Thermo Scientific, R904012) was melted at 65 °C before use. Then, 350 μL of Histogel was applied to a Cryomold (Tissue Tek, 4566) and spread evenly to eliminate bubbles and ensure complete coverage of all areas. Using a cell scraper, two confluent organoid domes were transferred onto the Histogel layer. Subsequently, 500 μL of warm Histogel was overlaid. The Histogel “sandwich” was solidified on ice for 25 minutes, then transferred between two sponges in a plastic cassette. Cassettes containing the organoid–Histogel sandwiches or patient tissue samples (obtained from Usask Biobank) were immersed in 10% neutral buffered formalin and fixed for approximately 24 h RT. After fixation, samples were transferred through at least two changes of 70% ethanol, each lasting approximately 24 hours, followed by tissue processing. Sections (4-5 μm thick) were air dried overnight at 37°C, baked at 60°C for 1 h and deparaffinized in xylene and rehydrated through a graded series of ethanol to distilled water. The sections were then stained with Harris’s hematoxylin for 1 minute, followed by rinsing in tap water. Differentiation was performed in Acid Ethanol (0.5% HCl in 95% Ethanol), and the sections were blued in saturated aqueous carbonate, then soaked in running tap water. Subsequently, the sections were stained with eosin for 2 minutes and then dehydrated through a graded ethanol series. They were cleared in xylene and mounted with a synthetic resin.

### Immunofluorescence of organoid cultures

CGM was aspirated from the 10 pancreatic organoid culture (log phase, passage numbers were among 5-8) and domes gently rinsed in PBS before fixing in 4% formaldehyde for 15 min. Domes were then permeabilized with blocking solution (0.1% Triton X-100, 3% BSA, in PBS) for 30 min at room temperature (RT). Domes were incubated with anti-EPCAM (CST, 2929S) and anti-E-Cadherin (CST, 3195S) primary antibodies diluted 1:1000 in blocking solution overnight at 4 °C Domes were then washed 3 x 10 min in wash buffer (0.1% Triton X-100 in PBS) and incubated with anti-mouse-Alexa-594 and anti-rabbit-Alexa-488 secondary antibodies diluted 1:1000 in blocking solution for 90 min at room temperature (Invitrogen, A11008; Invitrogen, A11005). Domes were subsequently washed 3 times for 10 min in wash buffer, counterstained with DAPI, and mounted (Thermo Fisher, P36961) onto glass slides for confocal imaging. All imaging was performed on a Zeiss LSM900 microscope.

### Whole Exome Sequencing (WES) on PDOs and corresponding patient samples

10 pancreatic organoid culture (log phase, passage numbers were among 5-8) pellets and 10 corresponding original patient samples (obtained from Usask Biobank) were sent to The Centre for Applied Genomics (TCAG) at the Hospital for Sick Children, Toronto, Canada (SickKids) for DNA extraction, quantification, quality control, library preparation and Whole Exome Sequencing. The exome capture was performed using the SureSelect Human All Exon V8 bait library (Agilent Technologies, 51918149) with the SureSelectXT HS2 Reagent Kit (Agilent Technologies, G9621A/G9622A). The final library was sequenced to a depth of 500x on one lane of a 25B flow cell using an Illumina NovaSeq 6000 with an S4 Reagent Kit (300 cycles) (Illumina, 20028313).

### Computational workflow for WES Data

Paired-end whole-exome sequencing reads from tumor tissues and matched organoid samples for each PANC samples were processed using a standardized computational pipeline. Raw FASTQ files were quality-filtered and adapter-trimmed using fastp (version 0.24.0) with automatic adapter trimming. High-quality reads were aligned to the human reference genome (hg38) using BWA-MEM (version 0.7.18). Alignments were sorted and indexed with samtools (version 1.18) and group information was added or corrected using GATK AddOrReplaceReadGroups to ensure compatibility with downstream variant calling. Somatic variant detection was performed using GATK Mutect2 (version 4.6.1.0), separately for tumor and organoid BAM files, using the GRCh38 reference genome, a population-germline resource (gnomAD), a panel of normals (PON), and a targeted interval file for the exome capture. Variants were filtered with GATK FilterMutectCalls to remove low-confidence calls.

Filtered VCFs were parsed into structured tables containing genomic coordinates, reference and alternate alleles, allele depth (AD), total read depth (DP), and variant allele frequency (VAF). Only variants with DP ≥ 12 and allele frequency ≥ 0.05 were retained for downstream analyses, ensuring sufficient read support for confident variant calls. Functional annotations from snpEff (ANN, which lists predicted effects for each alternate allele) or VEP (CSQ, which provides consequence annotations per transcript) were used to assign genes and predict consequences. Variants lacking gene annotations were intersected with exon coordinates using PyRanges and a reference BED file. Unique variant identifiers merged tumor tissue and organoid variant tables. Shared and sample-specific variants were identified based on VAF > 0. Gene-level analyses were performed by collapsing variants by gene and determining whether each gene harbored at least one mutation per sample. Mutational overlap and concordance were visualized using VAF scatter plots, gene-level percentage bar plots, and comparisons of mutational spectra. All computational and statistical visualizations were generated using Python libraries, including matplotlib, pandas, NumPy, and seaborn (version 0.13.2).

Somatic variant calls generated by GATK FilterMutectCalls were annotated using the Ensembl Variant Effect Predictor (VEP) web-based platform [43]. The Ensembl/GENCODE transcript database was used for transcript annotation, and allele-frequency data from the 1000 Genomes and gnomAD datasets were used to flag common population variants. The option to right-align variants prior to consequence calculation was enabled to ensure consistent indel representation. The annotated results were exported as a text file for downstream processing. To identify functionally relevant mutated genes associated with specific drug families in each organoid sample, an AWK-based parsing script was used to extract coding-region variants classified as *missense_variant, frameshift_variant, stop_gained, stop_lost, start_lost, splice_acceptor_variant, splice_donor_variant, inframe_insertion,* or *inframe_deletion.* The resulting non-redundant list of mutated genes was used to generate the heatmap shown in Figure 6.

### High-throughput *in vitro* drug sensitivity assay and enrichment analysis

A high-throughput *in vitro* drug screen was conducted using an FDA-approved drug library (TargetMol, L4200) containing 1,813 compounds, of which 61% were FDA-approved, and 38% were clinically used outside the United States. The screen was performed on 10 established pancreatic organoids (in log phase). Briefly, cells were dissociated with TrypLE (Gibco, 12605010) and seeded into Corning™ Matrigel™ Matrix-3D plates (Corning, 08-774-410) at a density of 3,000 cells per well in a volume of 50 µL using a Biomek NX^p^ liquid handling robot. Then, 200 nL of each drug compound from 500 µM stock plates (in DMSO) was transferred into each well using a drug pinning tool (V&P Scientific, BMPFXGR2PPER) operated by the robot, yielding a final drug concentration of 2 µM. After 96 hours of drug exposure, cell viability was assessed using the CellTiter-Glo® 3D Cell Viability Assay (Promega, G9683) according to the manufacturer’s instructions. Drug sensitivity was quantified using B-scores, which correct for positional and systematic plate effects using a median-polish procedure followed by variance scaling based on the median absolute deviation. B-scores were z-transformed across ten pancreatic cancer PDOs for cross-sample comparison, and compounds with normalized B-scores ≤ –2 were classified as inhibitory hits.

Compounds were categorized into primary pharmacological families using established drug knowledge bases supplemented with curated mechanism-of-action information from the literature. To identify shared vulnerabilities, drug-family enrichment was evaluated by comparing the number and weighted magnitude of inhibitory compounds within each family to expectations from a random distribution across the full screen (Fisher’s exact test). To integrate both potency and frequency of inhibition, a Weighted Enrichment Ratio (WER) was calculated:

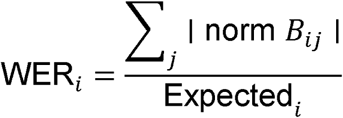

where | norm *B_ij_*| is the absolute normalized B-score for compound *j* in family *i*, and Expected is the mean weighted inhibition from random sampling. Families represented by ≥ 3 compounds were included, and significance values were adjusted using the Benjamini–Hochberg FDR procedure. Ranked WER and –log₁₀(FDR) values are shown in Figure 6A.

To profile inter-PDO pharmacologic heterogeneity, mean normalized inhibition values were calculated per active drug family for each PDO and visualized as a radar chart (Figure 6B). Overlapping regions represent shared drug sensitivities, whereas divergent polygon shapes denote PDO-specific pharmacologic fingerprints.

### Network analysis and mapping of PDO responses to standard-of-care drugs

A bipartite drug–organoid pharmacological network was generated in Cytoscape (version 3.10.4) to visualize shared and selective drug sensitivities across PDOs. Compounds classified as inhibitory hits (normalized B-score ≤ –2 in ≥1 PDO) were retained as drug nodes and annotated with primary pharmacological families. PDOs were represented as separate nodes, and edges were drawn between a PDO and a compound when the inhibitory threshold was met for that pair. A force-directed layout was applied to position compounds active in multiple PDOs centrally, while PDO-specific hits localized peripherally. A categorical drug response matrix was generated to compare PDO-specific sensitivity patterns across major chemotherapeutic and targeted drug classes. For each compound-PDO pair, normalized B-scores were categorized as Sensitive (≤ –2), Intermediate (–2 < B-score < 0), or Resistant (≥ 0). Compounds were grouped by clinical or mechanistic drug class, and responses were visualized in a bubble plot, where bubble color indicates the categorical response (green = sensitive, yellow = intermediate, red = resistant) and bubble size reflects the magnitude of inhibition. PDOs were ordered by hierarchical similarity, and drugs were arranged to highlight clinically relevant therapeutic categories in pancreatic cancer. This approach provides a comparative overview of pharmacologic heterogeneity across PDOs, revealing both shared and divergent drug responses.

## Acknowledgments

We thank the members of the Vizeacoumar and Freywald laboratories for their insights and comments on the manuscript. Banking was done by the University of Saskatchewan Biobank, founded by Ovarian Cancer Canada and supported by the University of Saskatchewan Pathology and Laboratory Medicine, which is affiliated with the Canadian Tumor Repository Network (CTRNet). This work was supported by a Saskatchewan Cancer Agency Operating grant, funded by donations to the Cancer Foundation of Saskatchewan. The Research Ethics Board of the University of Saskatchewan approved the study, in accordance with the appropriate regulatory authorities (USask Bio-REB 3382).

## Financial support

Canadian Institutes of Health Research operating grants PJT-156309, PJT-156401, and PLL-192134 (FJV, AF). Cancer Research Society operating funds 2017-OG-22493 (FJV, AF). Be Like Bruce Foundation (FJV, AF). Saskatchewan Cancer Agency operating grants from Cancer Foundation of Saskatchewan (AZ). College of Medicine, University of Saskatchewan (FSV). University of Saskatchewan CoMGRAD Award (VM). NSERC USRA (JQ), Canadian Institutes of Health Research - Canada Graduate Scholarships – Master’s (JDWP). University of Saskatchewan Health Science Graduate Scholarship (JDWP). Cancer Research Society – Doctoral Research Award (JDWP). Canadian Institutes of Health Research - Canada Graduate Scholarships – Doctoral FBD-187665 (VM).

## Author contributions

Conceptualization: F.S.V., A.F., F.J.V.

Investigation: H. D., F. S. V., Y. Z., N. J., J. D.W. P., V. M., L. G., T. F., J. P. V., M. L.- W., S. U., J. Q., A. M. M., A. R., R. K., Y. W., A. K., K. F., B. M., L. H., G. G., J. S., G. B., Y. L., M. O., K.B., M. M., S. A., A. Z.

Writing, reviewing, and editing: All authors

Funding Acquisition: A.Z., S.A., A.F., F.J.V.

## Declaration of interests

The authors declare no competing interests.

## Figure Legends

**Supplementary Figure 1.**
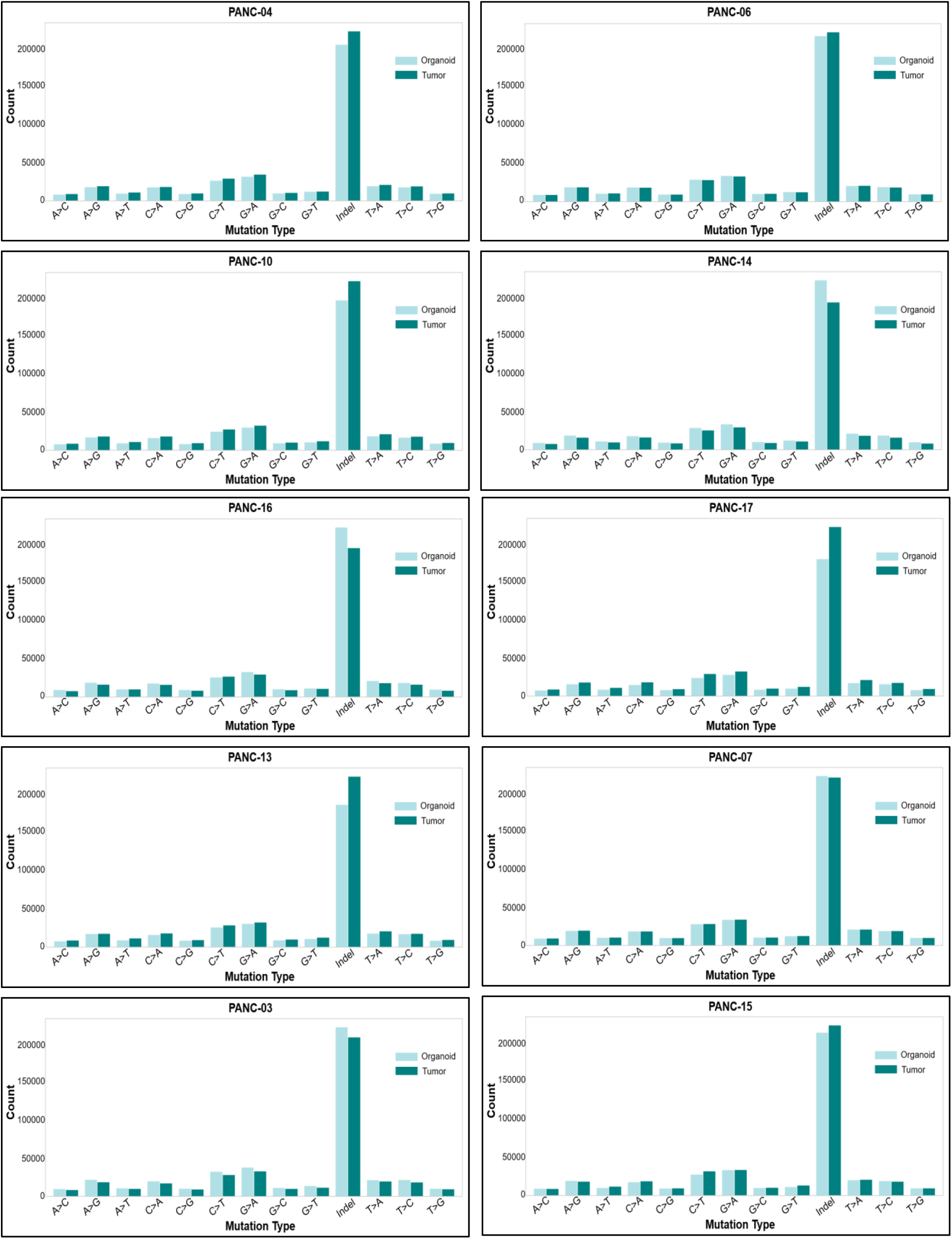
Mutational spectrum analysis of organoid–tumor pairs reveals conserved base substitution profiles across all PDO models. Bar plots display the distribution of single-nucleotide variants in each PDO model compared to its matched tumor, stratified by mutation type. Mutations are categorized into six base substitution classes (C>A, C>G, C>T, T>A, T>C, T>G), including their trinucleotide sequence contexts. Each panel represents a distinct PDO model–tumor pair (n = 10). Bars show mutation counts in the tumor (blue) versus the corresponding organoid model (green). Importantly, PDO models faithfully recapitulated the mutation spectrum of their parental tumors.

**Supplementary Figure 2.**
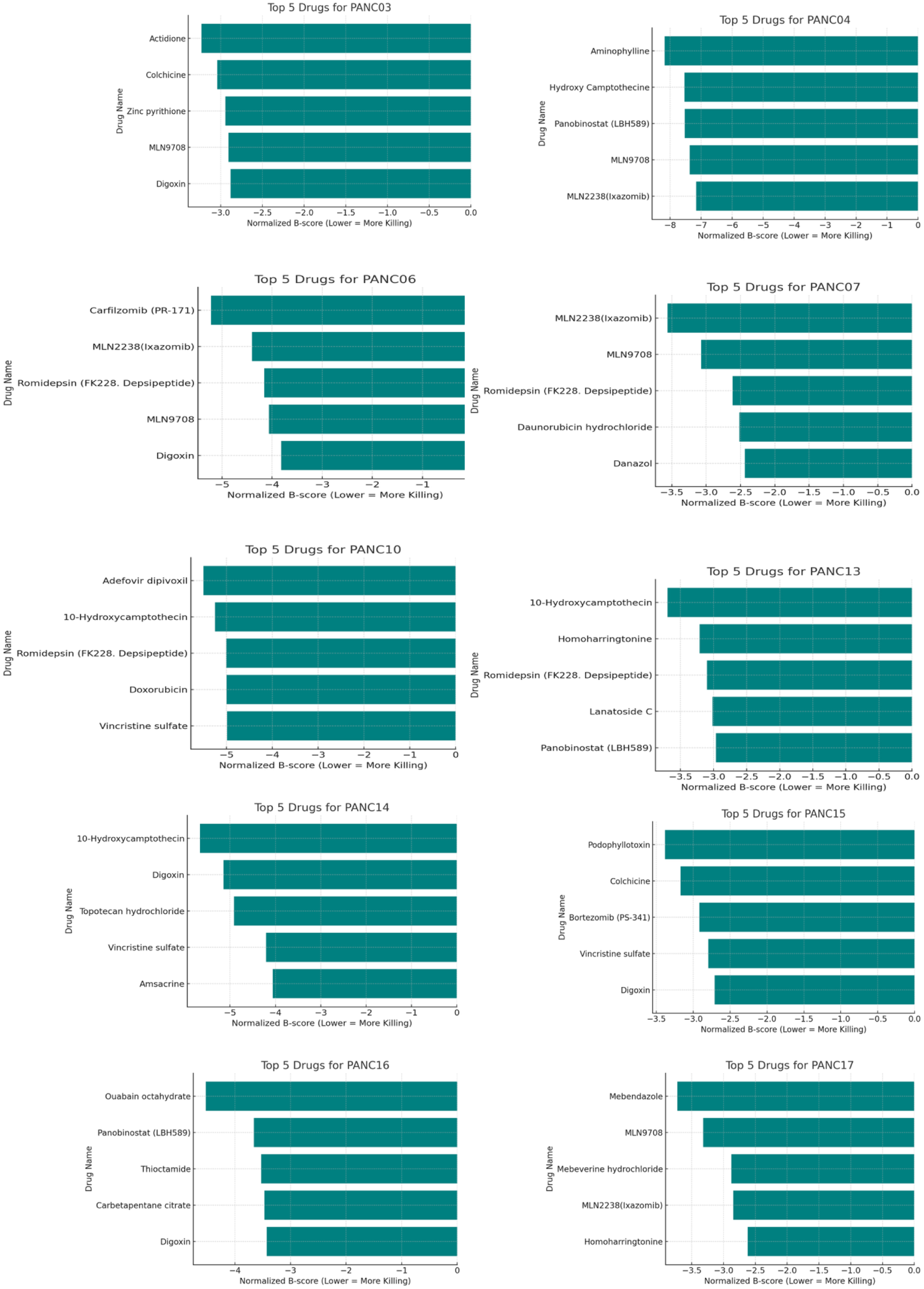
Top drug hits in each PDO model. For each of 10 PDO models (panels labeled by PANC ID), the bar graph shows five compounds with the lowest normalized B-score (highest efficacy). Bar length indicates the normalized B-score (with lower values indicating stronger killing). These plots identify the most active drugs for each PDO model, demonstrating that each PDO model has a distinct sensitivity profile.

**Supplementary Figure 3.**
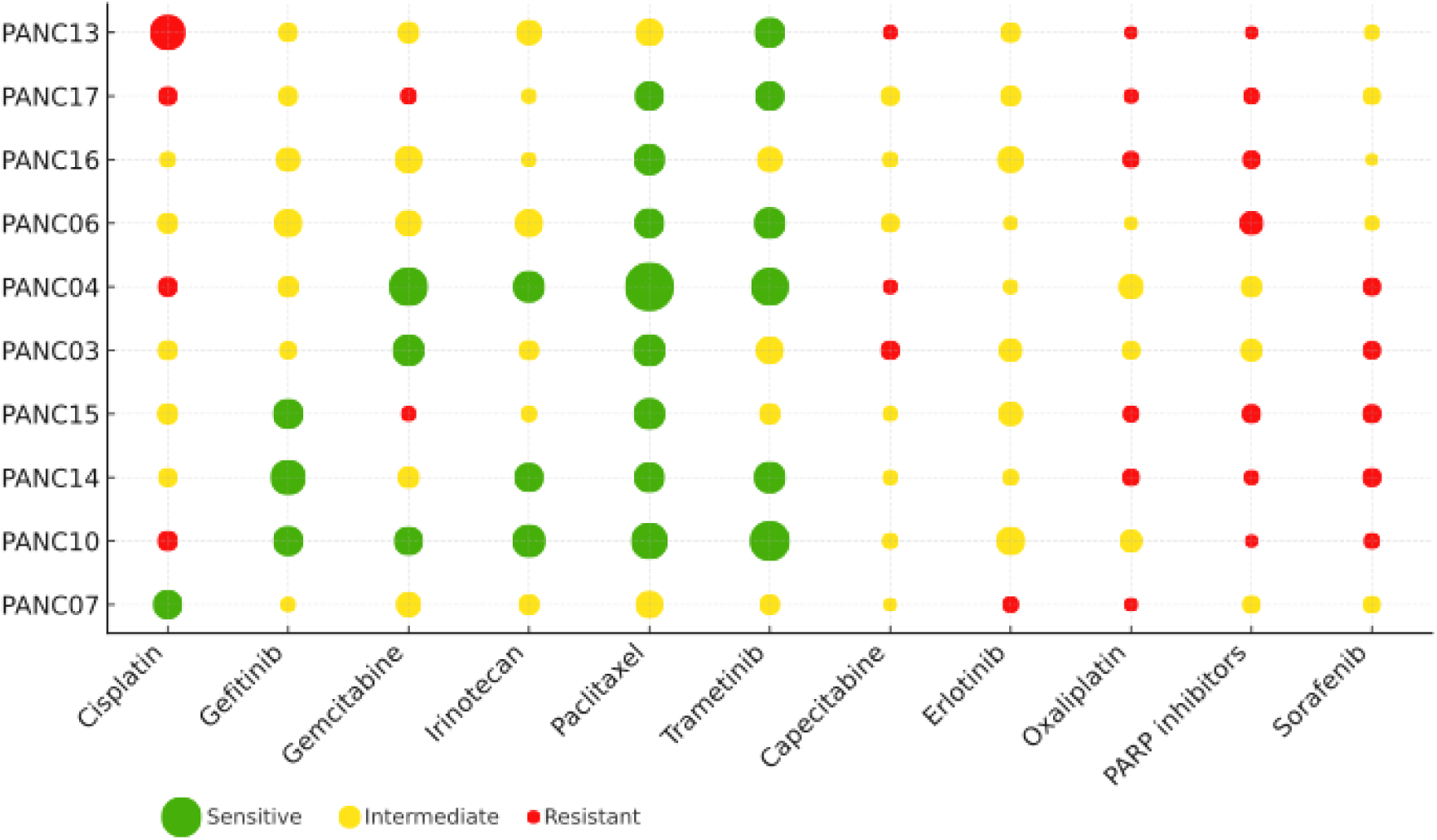
*Ex vivo* drug response profiles of PDO models to standard-of-care and investigational therapies. Bubble plot showing relative drug sensitivity of each PDO model to a panel of 12 clinically relevant agents, including standard chemotherapies (e.g., gemcitabine, paclitaxel, irinotecan), targeted agents (e.g., PARP inhibitors, trametinib, erlotinib), and repurposed compounds (e.g., cardiac glycosides). Each column represents individual drugs, and each row corresponds to a PDO model. The size of each bubble reflects the magnitude of the response, and the color indicates the response classification: green (sensitive), yellow (intermediate), or red (resistant), based on normalized B-score thresholds. While several PDOs showed broad sensitivity to agents like paclitaxel and gemcitabine, others exhibited selective resistance, highlighting intertumoral heterogeneity and the potential clinical value of PDO-guided therapy prioritization.

**Supplementary Figure 4.**
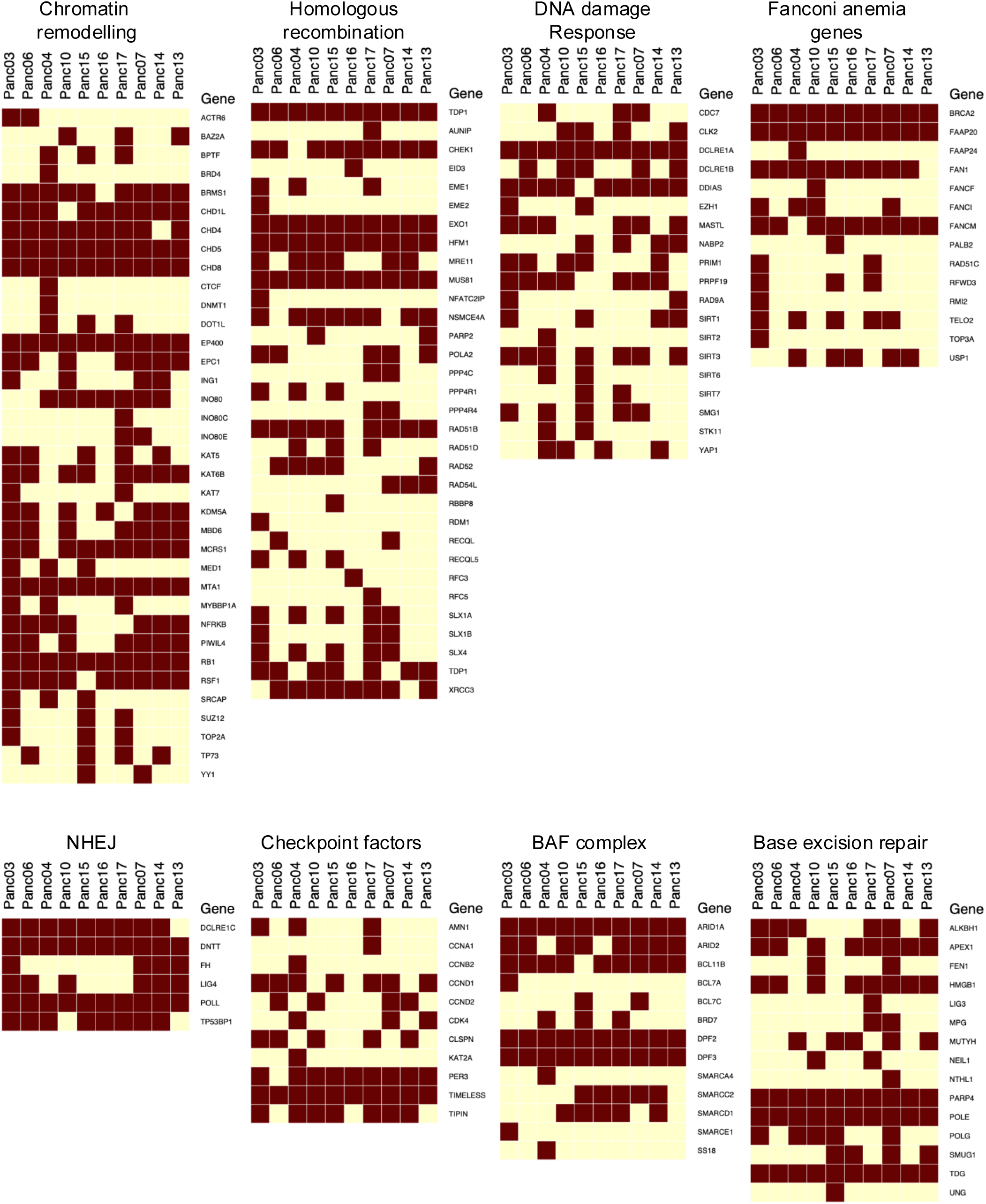
Expanded mutation profiling of DNA repair and chromatin-regulatory pathways across PDAC PDOs. Heatmaps displaying the presence or absence of somatic mutations across multiple DNA repair and chromatin-remodeling pathways in all ten PDAC PDO models. Each column represents a PDO, and each row corresponds to a gene within a specific functional category. Dark squares indicate detected mutations; pale squares indicate wild-type status.

## Notes

### Competing Interest Statement

The authors have declared no competing interest.

